# DNA replication initiation drives focal mutagenesis and rearrangements in human cancers

**DOI:** 10.1101/2024.03.12.584624

**Authors:** Pierre Murat, Guillaume Guilbaud, Julian E. Sale

## Abstract

The rate and pattern of mutagenesis in cancer genomes is significantly influenced by DNA accessibility and active biological processes. Here we show that efficient sites of replication initiation drive and modulate specific mutational processes in cancer. Sites of replication initiation impede nucleotide excision repair in melanoma and are off-targets for activation-induced deaminase (AICDA) activity in lymphomas. Using ductal pancreatic adenocarcinoma as a cancer model, we demonstrate that the initiation of DNA synthesis is error-prone at G-quadruplex-forming sequences in tumours displaying markers of replication stress, resulting in a previously recognised but uncharacterised mutational signature. Finally, we demonstrate that replication origins serve as hotspots for genomic rearrangements, including structural and copy number variations. These findings reveal replication origins as functional regulators of tumour biology and demonstrate that replication initiation both passively and actively drives focal mutagenesis in cancer genomes.

## Introduction

Cancer cells are characterised by their ability to sustain cell growth and proliferation by evading suppressor signals and activating oncogene expression. ^1^ The overexpression of oncogenes such as c-Myc, h-RAS, or Cyclin E shortens the G1 phase of the cell cycle, expedites S phase entry, and increases levels of replication initiation. ^2,3,4^ This accelerated S-phase entry is linked to replication stress and genome instability, in part due to origin over-usage and activation of dormant replication origins. ^5^ Recent work has shown replication origins are hotspots for mutagenesis in the germline. ^6^ However, the impact of replication initiation on mutagenesis in human cancers has not been examined.

We recently reported that a subset of highly efficient replication origins, conserved across a range of untransformed, immortalised and transformed human cell lines, act as hotspots for mutagenesis in the germline. ^6^ These ‘constitutive’ replication origins are notably enriched within the active and early replicated regions of the human genome, particularly within gene bodies. The mutational processes operating at these sites thus contribute to elevated mutational burdens at gene promoters and splice junctions, impacting phenotypic diversity across human populations. Origin activity has also been shown to drive translocations at the *MYC* locus in murine B cells, suggesting a potential link between replication origins and the development of lymphomas and other cancers. ^7^ Further, increased origin firing has been associated with chromosomal instability in human colorectal cancer cell lines, with overexpression of origin firing genes such as GINS1 and CDC45 resulting in aneuploidy. ^8^ Collectively, these studies highlight replication origin firing as a cancer-relevant trigger for mutagenesis and genome rearrangements.

In this study, we report the identification and characterisation of mutational processes occurring at constitutive origins in cancer. Our results emphasise the major impact of replication initiation in shaping the mutational landscape of the human genome, adding replication origins to the expanding repertoire of genomic features influencing local mutation rates in cancer (reviewed in ^9^).

## Results

### Pan-cancer analysis of mutation distribution reveals enrichment at replication origins

To systematically examine the frequency of somatic point mutations at constitutive origins (refer to the **Online Methods** section for detailed information on the analysed origin sites), we considered mutation calls from the International Cancer Genome Consortium (ICGC) aggregated from 86 cancer projects. We found a ∼ 1.7-fold enrichment in single nucleotide variants, hereafter refer to as mutations, at these sites compared to their flanks (**Extended Data Fig. 1a**). This was accompanied by an increase in minor allele frequencies (**Extended Data Fig. 1b**). To account for local variation in base composition, we corrected the apparent mutation rates of each of the six pyrimidines mutation by their occurrences and found increased mutation rates associated to each type (**Fig. 1a**). These observations suggest that a specific form of mutagenesis is focused at constitutive origins in cancer genomes.

**Figure 1.**
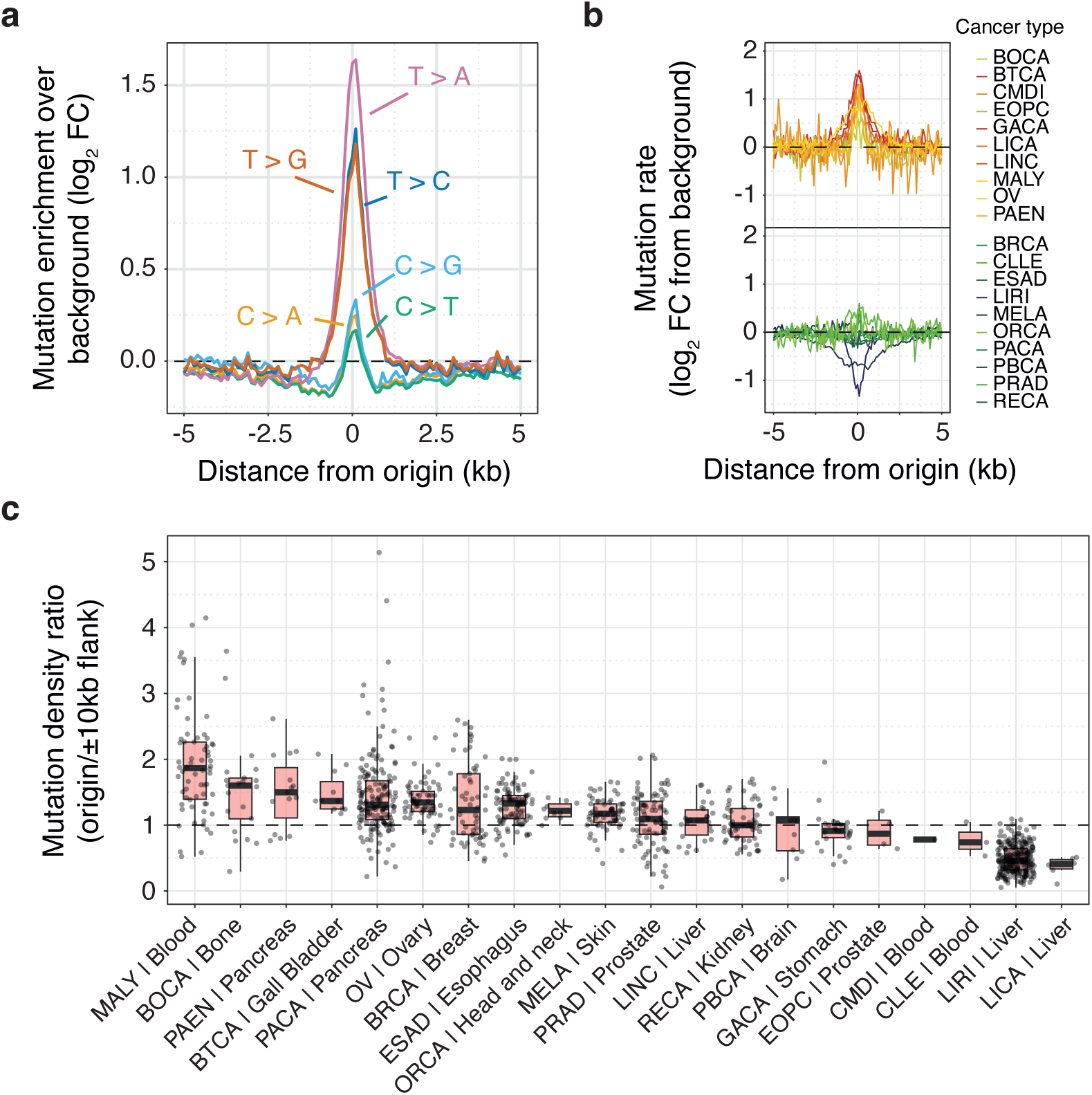
Pan-cancer analysis of mutation distribution across the genome reveals enrichment at constitutive origins. **a**, Mutation rates associated with the six pyrimidine substitutions at constitutive origins. Mutation rates were computed from aggregated mutation calls from 86 ICGC cancer projects, corrected for local variation in base composition and background values. **b,** Averaged and background adjusted mutation rates stratified by cancer types. Acronyms of cancer types are detailed in the **Online Methods** section. **c**, Distribution of origins / origin flanks mutation density ratios of individual cancer samples stratified by cancer type and primary sites. Density ratios were computed by considering origin domains (origin midpoints ± 500 bp) and origin flanking domains (origin midpoints ± 10 kb excluding origin domains). Box plots show medians and interquartile ranges. Individual cancer samples are shown as grey dots. Only whole-genome-sequenced cancer samples with at least 5,000 mutations were considered for experiments reported in panels **b** and **c**.

We then asked whether mutation rates at constitutive origins vary across cancer types. We computed origin mutation rates for individual tumours with over 5,000 mutations (1,056 tumours across 20 cancer types) to ensure reliable estimates of mutation density at and near origins. While some cancers, like gastric and gall bladder, show high mutation rates at origins (**Fig. 1b**), others, like renal or liver cancers, unexpectedly have constant or even negative rates. To assess variation within cancer types, we computed the ratios of mutations at origins compared to flanking regions and assessed their variability across cancer types (**Fig. 1c**). This revealed significant differences within specific cancer types, pancreatic cancer, for instance, showing ratios ranging from 0.21 to 5.14. These observations suggest that increased mutation rates at origins are driven by mutational processes of varying degrees of intensity rather than inherent tumour properties. Supporting this, we found a strong correlation between the total number of mutations and those at origins (*Rho* = 0.954, **Extended Data Fig. 1c**), but an opposite correlation with the origin/flank mutation ratio (*Rho* = −0.246, **Extended Data Fig. 1d**). This pattern suggests that specific mutational processes target constitutive origins.

### Origin-associated mutational signatures

To identify and characterise these mutational processes, we clustered cancer samples based on the cosine similarity of their origin trinucleotide mutation signature (**Fig. 2a**). Our analysis specifically focused on tumours with over 50 mutations at origins (227 tumours across 17 cancer types) to ensure sufficient power to call distinct signatures. Each origin signature underwent adjustment for local trinucleotide composition and correction to account for background values from the flanking domains (details in **Online Methods**). The clustering revealed five distinct tumour groups, with two clusters exclusive to a specific cancer type and three including multiple types (**Fig. 2b**). Subsequently, representative mutation signatures were computed by aggregating mutations at origins from tumours within each cluster (**Fig. 2c**). These were compared to known signatures of somatic mutations in cancerous human tissues (COSMIC SBS signatures, **Extended Data Fig. 2a**). ^10^

**Figure 2.**
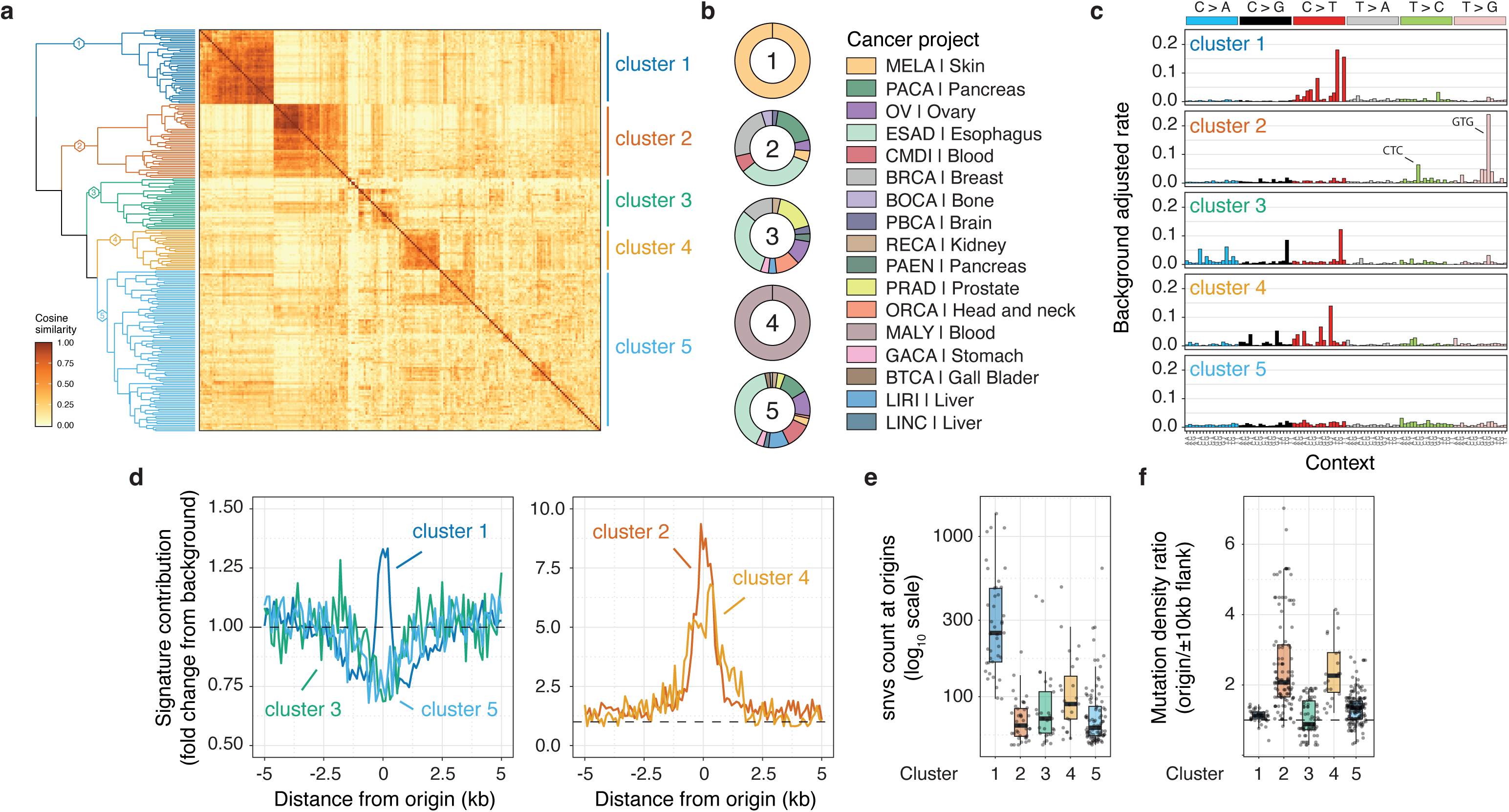
Signature analysis uncovers distinct mutagenic processes focused at constitutive origins. **a**, Cancer sample clustering based on the cosine similarity of their origin trinucleotide mutational signature. Each signature at origin domains was corrected to adjust for local trinucleotide composition and background values from neighbouring origin flanking domains. Only whole-genome-sequenced cancer samples with at least 50 mutations at origins were considered. **b**, Distribution of cancer types and primary sites within the five identified clusters of tumours. **c**, Origin-associated mutational signatures for tumour clusters. Signatures were computed from aggregated mutation calls at origin domains for clustered cancer samples and corrected as previously. **d**, Local exposure to mutational signatures associated with clustered tumours. Signature contributions are corrected for background origin flank values. **e**, Total mutation count at origins and (**f**) origin/origin flanks mutation density ratios for individual cancer samples grouped by clusters. Box plot show medians and interquartile ranges. Individual cancer samples are represented as grey dots.

Clusters 1 and 4 are specific to skin melanomas (MELA) and malignant lymphomas (MALY) respectively, displaying characteristic origin signatures highly similar to the known signatures SBS7b (*cosine similarity* = 0.83, exposure to ultraviolet light) and SBS84 (*cosine similarity* = 0.88, activity of activation-induced cytidine deaminase (AID or AICDA)). Cluster 2 encompasses tumours from various cancer types, with pancreatic ductal adenocarcinoma (PACA), oesophageal adenocarcinoma (ESAD), and breast cancer (BRCA) being the primary contributors. Cluster 2 signature exhibits high similarity to SBS43 (*cosine similarity* = 0.95), a signature of unknown aetiology, marked by T > G mutations in the specific context GTG. Clusters 3 and 5 comprise samples from over 10 different cancer types. Cluster 3 signature shares moderate similarities with the known SBS10b signature (*cosine similarity* = 0.64, associated with polymerase epsilon exonuclease domain mutations). Lastly, the aggregated cluster 5 signature appears featureless, resembling a ‘flat’ signature probably resulting from merging signals originating from diverse processes. It is noteworthy that clusters 3 and 5 are the least resolved clusters in our analysis (**Fig. 2a**). We then compared these origin-associated signatures with genome-wide signatures derived from samples within the respective clusters (**Extended Data Fig. 2b**). Aside from cluster 1, our origin-associated signatures exhibit minimal resemblance to the corresponding genome-wide signatures. This finding validates our approach for identifying processes that focus mutagenesis at origins.

To gain a deeper understanding of the specificity of our origin-associated signatures, we examined their local contribution near origins (**Fig. 2d**). The aggregate origin signatures derived for clusters 3 and 5 show no discernible contribution to origin mutagenesis. Conversely, processes linked to cluster 2 and 4 signatures focus mutations at origins, with a significant 7- to 9-fold increase compared to origin flanks. The process related to the cluster 1 signature exhibits some preference for origins, albeit with a moderate 1.3-fold enrichment. We further quantified the number of mutations attributed to each signature at origins (**Fig. 2e**) and evaluated the specificity of associated processes (**Fig. 2f**) at the level of individual cancer samples. These analyses reveal that while the process at origins within cluster 1 lacks specificity, it significantly contributes to origin mutagenesis. Conversely, clusters 2 and 4 are highly specific to origins but contribute moderately in terms of absolute mutation numbers. We validated these findings by fitting COSMIC SBS signatures to mutational profiles of origin domains (**Extended Data Fig. 2c**). Overall, our analysis identifies three independent mutational processes targeting mutagenesis at constitutive origins, two of which are tissue-specific, while one operates across different cancer types. Next, our aim was to identify the aetiology of these processes, beginning with those specific to particular tissues.

### Differential DNA repair underlies mutation hotspots at origins in skin melanoma

Samples from cluster 1 exclusively originate from skin melanomas and exhibit both origin and genome-wide signatures that closely resembles the known COSMIC SBS7b (**Fig. 2b-c** and **Extended Data Fig. 2b**). However, the absolute contribution of SBS7b is focused at origins with approximately 2.5-fold higher values at origins compared to their flanks (**Extended Data Fig. 2c**). This could be explained by either locally impaired DNA repair or a higher probability of UV-induced lesions at constitutive origins. Given that both protein binding and transcription initiation have been reported to impair nucleotide excision repair (NER), ^11,12^ we hypothesised that increased mutagenesis at origins may result from NER deficiency due to the loading of the pre-replication complex at licensed origins.

To assess this point, we used genome-wide maps of NER activity, obtained through XR-seq analysis of irradiated skin fibroblasts for cyclobutane pyrimidine dimers (CPD) and pyrimidine-pyrimidone (6-4) photoproducts (6-4 PP). ^13^ We correlated the XR-seq profiles with DNase I hypersensitivity (DHS) at origins (**Extended Data Fig. 3a**) and observed a positive correlation between NER activity and origin accessibility (*Rho* = 0.40 and 0.17 for CPD and 6-4 PP respectively, **Extended Data Fig. 3b**) suggesting increased accessibility is associated with higher levels of damage. However, we found that the profile of mutation density at origins inversely mirrors NER activity (**Fig. 3a** and **Extended Data Fig. 3c**) and that the number of mutations at origins displayed a negative correlation with XR-seq signals at these sites (*Rho* = −0.44 and −0.36 for CPD and 6-4 PP respectively, **Fig. 3b**). Thus, replication initiation may specifically impair NER explaining the increased in SBS7b mutagenesis at origins in melanoma. It worth noting that we performed this analysis using data from CSB/ERCC6 mutants to interrogate global NER rather transcription-coupled NER. We further ruled out the involvement of transcription-coupled NER at origins by observing that XR-seq profiles at intergenic origins were comparable to those observed for origins located at promoters or within gene bodies (**Extended Data Fig. 3d**).

**Figure 3.**
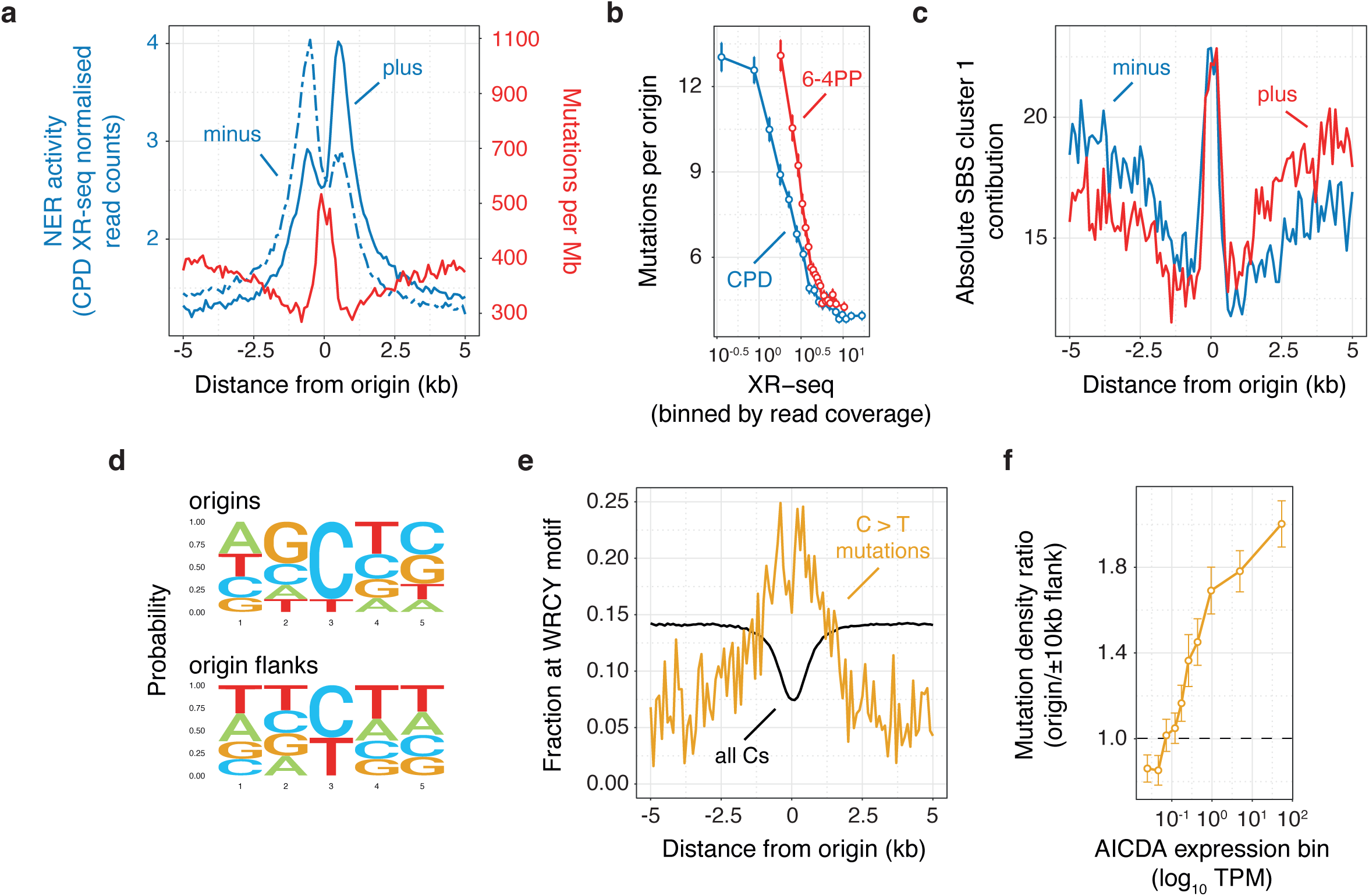
Differential DNA repair and off-target AID deamination underlie mutation hotspots at origins. Nucleotide excision repair in ultraviolet-irradiated human cells inversely mirrors mutation density at origins in skin melanomas. **a**, Average melanoma mutation density for cluster 1 tumour samples (red line) and strand-resolved XR-seq profiles for CPD (blue lines) in CSB/ERCC6 mutant NHF1 skin fibroblasts at constitutive origins. **b**, Number of mutations at origins as a function of XR-seq signals for CPD (blue line) or 6-4 PP (red line). XR-seq signals were binned by read coverage. Mutation data represent the means and standard errors to the mean of values for each XR-seq signal bin. **c**, Strand-resolved cluster 1 mutational signature contribution to mutagenesis at the origins of cluster 1 tumours. Signature contribution was computed by considering aggregated mutation calls from cluster 1 tumours. Constitutive origins are off-targets for AID deamination in malignant lymphomas. **d**, Consensus contexts of mutations mapped within origin or origin flank domains for cluster 4 tumours. The central position represents the mutated bases. **e**, Fraction of C > T mutations (orange line) or C residues (black line) overlapping the WRCY AID hotspot motif (where W represents weak bases, R purines, C the mutated bases and Y pyrimidines) at origins of cluster 4 tumours. **f**, Pan-cancer origins/origin flanks mutation density ratios as a function of AID (AICDA) expression. Mutation density ratio values were computed for transcript per million (TPM) bins and represent the means and standard errors of the mean.

Interestingly, when examining strand-resolved XR-seq signals (**Fig. 3a** and **Extended Data Fig. 3c**), we observed a preference for NER activity on the lagging strand near constitutive origins. To determine whether this observation arises from a replication strand bias or from a skew in UV-reactive dinucleotides (such as TT dinucleotides), we adjusted XR-seq signals for dinucleotide composition skew (details in **Online Methods**) and calculated strand biases (**Extended Data Fig. 3e**). Despite the correction, NER activity still exhibited a bias toward the lagging strand. Subsequently, we evaluated the strand specificity of our cluster 1 origin-associated signature (**Fig. 3c**) and found no strand bias at the origins but an enhanced contribution on the lagging strand in close proximity to the origin. Taken together, these findings support NER deficiency at origins and increased NER on the lagging strand of replication forks translocating away from origins.

### Origins are off-targets of cytosine deaminases in lymphomas

The cluster 4 signature (**Fig. 2c**) exhibits notable specificity for origins (**Extended Data Fig. 2b**) and bears significant similarity to the established COSMIC SBS84 signature associated with AID activity in lymphoid cancers. We hypothesised that origins might serve as off-targets for AID deamination. To investigate this point, we first examined the extended context of cluster 4 mutations at origins or their flanks (**Extended Data Fig. 3f**). Our analysis revealed that C > T mutations at origins occur within specific extended contexts, distinct from those observed at origin flanks, characterised by an enrichment of As, Gs, and Cs at the −2, −1, and +1 positions relative to the C > T mutations. This mutation context closely resembles the known WRCY AID hotspot motif (where W represents weak bases, R purines, C the mutated bases and Y pyrimidines). ^14,15^ Subsequently, by constructing consensus sequences for C > T mutations at origins and origin flanks (**Fig. 3d**), we confirmed that mutagenesis at origins in Cluster 4 genomes aligns with this consensus sequence. Furthermore, to demonstrate the activity of AID at origins in lymphomas, we calculated the fraction of C > T mutations within the canonical WRCY motif around origins (**Fig. 3e**) and observed a local increase at origins. Interestingly, the fraction of Cs in a similar context decreased at origins, supporting that origins are indeed off-targets of AID. To reinforce this observation, we identified a pan-cancer correlation between mutational burden at origins and AID expression (*Rho* = 0.25, **Fig. 3f**).

Another source of deamination in lymphomas and other cancer genomes arises from the activity of APOBEC deaminases. ^16,17^ We therefore sought to determine whether APOBEC activity contributes to mutagenesis at origins. To investigate this, we initially examined whether high expression of APOBEC in lymphoma cancers is associated with increased mutational burden at origins (**Extended Data Fig. 3g**). Our analysis revealed that the overexpression of APOBEC3 enzymes is indeed associated with higher origin/flank mutation density ratios, suggesting a potential contribution of APOBEC3 to origin mutagenesis. However, when assessing the background-adjusted origin mutation profiles computed for lymphoma genomes, we observed minimal contribution from the COSMIC SBS2 and SBS13 signatures, which are attributed to the activity of the APOBEC family of cytidine deaminases (**Extended Data Fig. 3h**). This observation suggests that cytosine deamination at origins in lymphomas is primarily driven by AID.

### Polymerase δ behaviour at non-B DNA structures drives origin mutagenesis

Cluster 2 samples stems from a wide range of cancer types, with focal mutagenesis at origins primarily driven by T > G mutations (see **Fig. 2c** and **Extended Data Fig. 4a**). Investigating this distinctive signature, we first analysed the extended contexts of these mutations at origins and their flanks (**Fig. 4a**). Interestingly, T > G mutations at origins, unlike those within origin flanks, tend to occur within G-rich sequences, particularly embedded within G tracts. This suggests a potential association with non-B DNA structures, such as G-quadruplexes (G4s), known to hinder replication and induce mutagenesis and genetic instability. ^18,19^ To probe the role of G4s in mutagenesis at origins, we analysed short sequences overlapping with T > G mutations at origins and their flanks (25 bp around mutations), predicting their susceptibility to fold into G4 structures using the G4Hunter algorithm, which considers G-richness and G-skewness to generate a quadruplex propensity score (G4H score). ^20^ Our analysis reveals that sequences surrounding T > G mutations at origins in cluster 2 genomes exhibit significantly higher overall positive G4H scores compared to sequences containing T > G mutations at origin flanks or sequences containing any T residues at origins (**Fig. 4b**). This indicates a potential driving role of G4 structures in mutagenesis at origins. We next identified sequences with the potential for G4 formation following a N_5_G_3+_N_1–12_G_3+_N_1–12_G_3+_N_1–12_G_3+_N_5_ pattern (where N is any base) ^21^ within origin domains, and computed their G4H scores. Remarkably, sequences bearing T > G mutations display higher G4H scores than those without mutations (**Fig. 4c**), suggesting that G4 structures, rather than solely G-rich sequences, are driving mutagenesis, with the most stable predicted G4s being the most mutated.

**Figure 4.**
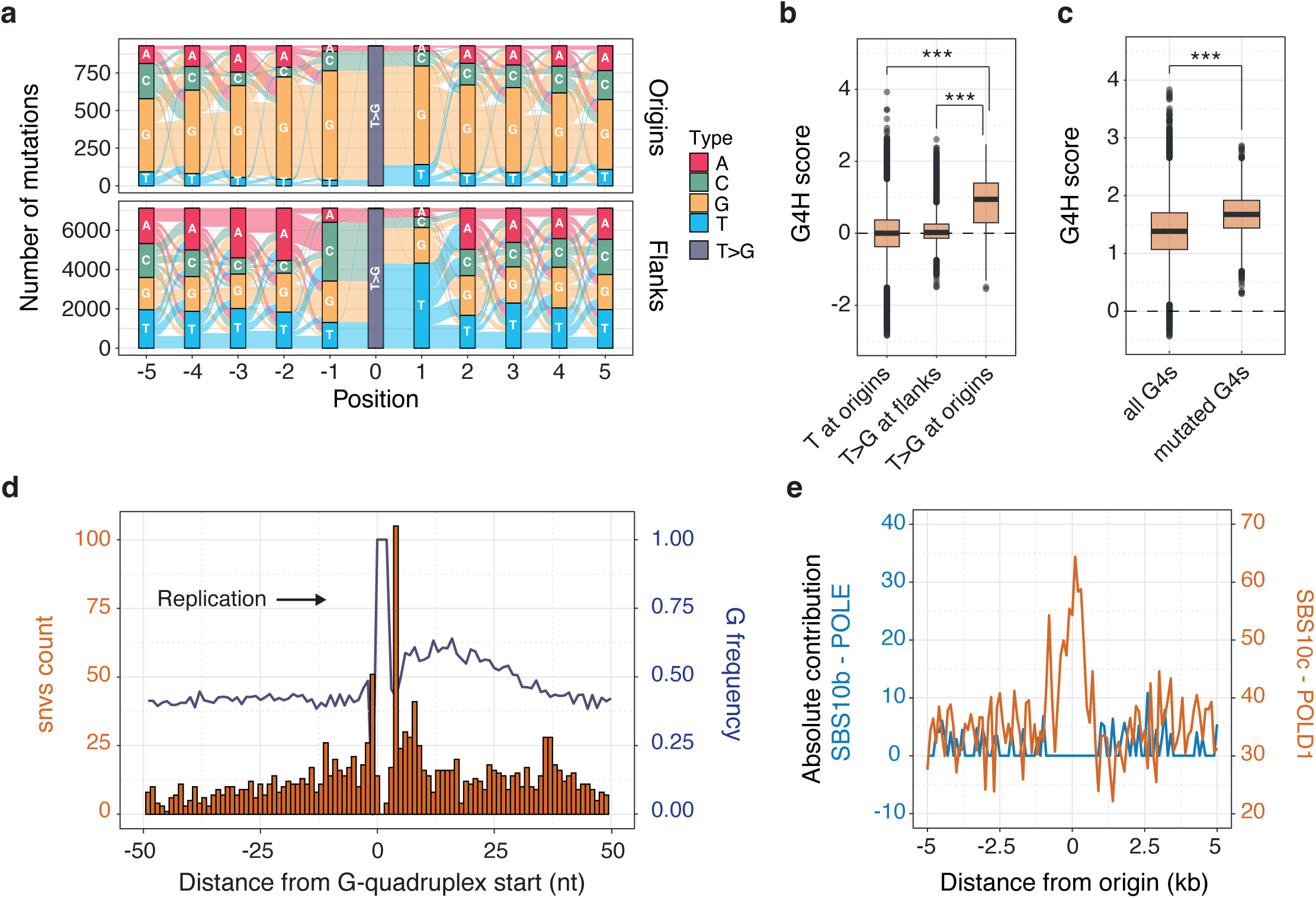
Polymerase δ behaviour at G-quadruplex structures drives origin mutagenesis in diverse cancer types. **a**, River plot illustrating the sequence contexts of T > G mutations at origins or origin flanks for cluster 2 tumour samples. **b**, Distribution of the G-quadruplex (G4) propensity scores, G4H scores, associated with short 51 bp sequences encompassing T > G substitutions at origins and origin flanks, or any T residues at origins. **c**, Distribution of G4H scores for all or T > G mutated sequences at origin domains, fitting the pattern N_5_G_3+_N_1– 12_G_3+_N_1–12_G_3+_N_1–12_G_3+_N_5_ (where N is any base), representing potential G4 formation. Box plot report medians and interquartile ranges. Outliers are shown as grey dots. \*\*\**P* < 0.001, Kolmogorov-Smirnov test. **d**, Distribution at single-nucleotide resolution of mutations and G frequency at G4-forming sequences found within 1 kb domains centred on origins. The start of the G4s is defined as the first G from the first G tract of the previous pattern. G4-forming sequences were oriented toward the direction of replication. **e**, Absolute contribution of COSMIC signatures SBS10b and SBS10c to mutagenesis at origins, associated with the activity of POLE and POLD1, respectively. Both signatures are attributed to defective proofreading due to acquired mutations in the exonuclease domains of the polymerases.

Having established the role of G4s in driving mutagenesis at origins, we delved deeper into understanding the characteristics of our cluster 2 origin signature. We hypothesised that T > G mutations in the specific GTG context (**Fig. 2c**) might be disproportionately represented in G4-forming sequences. We analysed the occurrence of each of the 96 trinucleotide contexts within the established G4 origin pattern and revealed an enrichment of the GTG context within G4 forming sequences at origins (**Extended Data Fig. 4b**). Accounting for the trinucleotide composition of G4s enabled us to extract a mutational signature specific to G4s (**Extended Data Fig. 4c**), illustrating the diverse spectrum of mutations observed at these sites. Furthermore, we examined the mutation pattern at origin G4s by precisely mapping mutations at G4 sites oriented by the direction of replication (**Fig. 4d**). Our findings indicate an accumulation of mutations at the first nucleotides before and after the initial G tract of G4s, aligning with biochemical observations, ^22,23^ suggesting that polymerases may stall or proceed with errors past the first G tract of G4s during polymerase extension assays. This supports that mutagenesis at origins in cluster 2 is primarily a result of polymerase stalling at G4s. However, we observed that our cluster 2 signature does not exhibit a replication strand bias at origins (**Extended Data Fig. 4d**), and that G4s at origins are not predicted to be inherently more stable than G4s at their flanks (**Extended Data Fig. 4e**). This suggests that a single replicative polymerase may be responsible for mutagenesis at origins, and its synthesis errors are confined within origin domains.

Evidence, from both biochemical studies in yeast and genetic investigations in humans, ^6,24,25,26^ have elucidated that following the priming of DNA replication by polymerase α, polymerase δ initiates both leading and lagging strand synthesis, thus performing the bulk of DNA synthesis near replication origins. This initial DNA synthesis following origin firing occurs uncoupled from helicase activity and is likely more susceptible to polymerase stalling, potentially accounting for the increased mutation rate observed at G4s near origins. Consequently, we evaluated the exposure of origins to mutational signatures associated with the activity of POLD1 and POLE, which become more pronounced when the exonuclease domains of these polymerases are compromised. ^27^ Our analysis revealed that the local distribution of these signatures aligns with polymerase δ being the active polymerase within 1-kilobase domains centred on human origins (**Fig. 4e**). Taken together, these findings support a model wherein polymerase δ stalls and triggers error-prone DNA synthesis at G4 structures in the immediate vicinity of replication origins of cluster 2 tumours.

### Mutagenesis at origins is exacerbated by increased proliferative drive and replicative stress

DNA secondary structure-dependent mutagenesis at origins is shared among various cancer types, although it is not uniformly observed within all samples of the same cancer type. Our next goal was to elucidate the specific cancer features that influence the intensity of mutagenesis at origins. We selected pancreatic ductal adenocarcinoma (PACA) as a model cancer due to its wide range of variation in the ratio of mutations at origins to flanks (**Fig. 1c**). We first identified distinct cancer subtypes within PACA based on transcriptomic profiles. Using a uniform manifold approximation and projection dimensionality reduction (UMAP) on normalised gene expression counts of individual samples (**Fig. 5a**), we identified three subtypes, henceforth referred to as PACA 1 to 3, each exhibiting different levels of mutagenesis at origins. While all subtypes show similar genome-wide mutation burdens (**Extended Data Fig. 5a**), PACA 1 exhibits a higher degree of mutagenesis at origins compared to PACA 2 and 3 (**Fig. 5b**). Notably, tumours classified as PACA 1 were characterised by the presence of our cluster 2 origin signature, a feature attenuated in PACA 2 and absent in PACA 3 (**Extended Data Fig. 5b**), suggesting that DNA structure-dependent mutagenesis at origins occurs to varying extents in these cancer subtypes. Consistently, exposure to our cluster 2 signature at origins was higher in PACA 1 compared to PACA 2 and 3 (**Fig. 5c**). These observations indicate that UMAP reduction enables the stratification of pancreatic cancers into distinct subtypes, facilitating the study of determinants influencing mutagenesis at origins.

**Figure 5.**
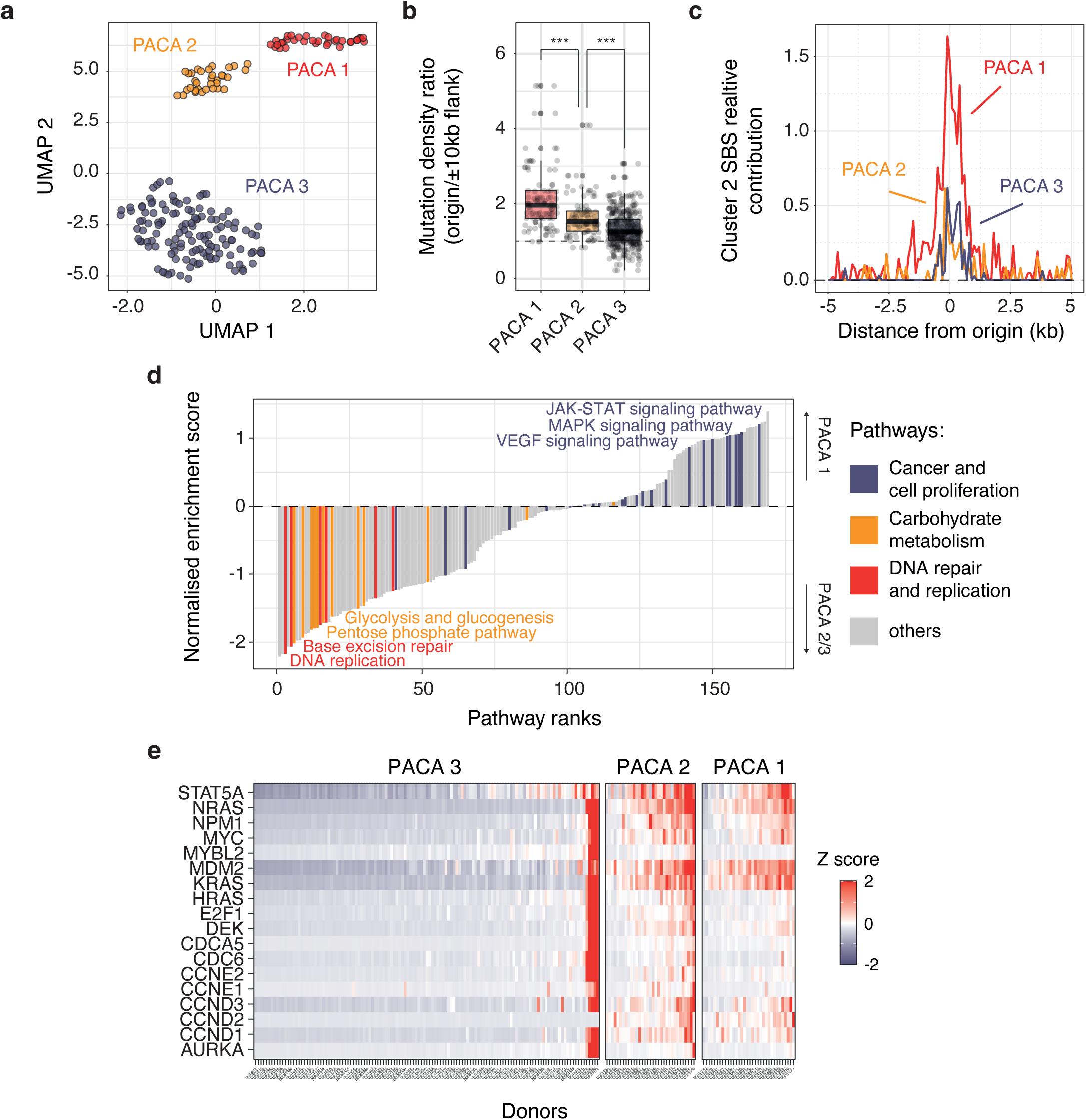
Replicative stress exacerbates origin mutagenesis in pancreatic ductal adenocarcinoma. **a**, UMAP dimensionality reduction of transcriptomic profiles from 194 pancreatic ductal adenocarcinoma samples (PACA project) reveals three cancer subtypes denoted PACA 1 to 3. **b**, Distribution of origins/origin flanks mutation density ratios for individual cancer samples categorised by PACA subtypes. Box plot report medians and interquartile ranges. Grey dots represent individual cancer samples. \*\*\**P* < 0.001, Kolmogorov-Smirnov test. **c**, Relative contribution of Cluster 2 signature to mutagenesis at origins. **d**, Pathway enrichment analysis of genes dysregulated in PACA subtype 1 compared to subtypes 2 and 3. Normalised enrichment scores indicate the extent of upregulation of gene sets representing KEGG pathways in PACA 1 tumours. Selected dysregulated pathways are colour-coded: cancer and cell proliferation (blue), carbohydrate metabolism (orange), and DNA repair and replication (red). **e**, Gene expression levels of replication stress biomarkers in pancreatic cancer, ^30^ categorised by PACA subtypes, represented as Z scores computed from the distribution of gene expression values across all PACA samples.

We proceeded with a differential gene expression analysis to identify genes displaying altered expression in PACA 1 compared to PACA 2 and 3 (**Extended Data Fig. 5c**). Subsequently, the results of this analysis were used for a gene set enrichment analysis (GSEA) to explore the altered KEGG pathways in PACA 1 (**Fig. 5d** and **Extended Data Fig. 5d**, details in **Online Methods**). The GSEA revealed the upregulation of cancer pathways (*P* = 7.44 × 10^-5^, permutation test) linked with increased cell growth and proliferation, such as the JAK-STAT or MAPK pathways. ^28,29^ This led us to hypothesise that increased mutation rates at origins might stem directly from replicative stress induced by excessive cell proliferation. To address this hypothesis, we evaluated the expression of biomarkers associated with replicative stress in pancreatic cancers ^30^ and discovered that tumours from PACA 1 and PACA 2 subtypes indeed exhibit cellular stresses interfering with DNA replication (**Fig. 5e**), contrasting with the absence of such stresses in PACA 3. However, PACA 1 and PACA 2 tumours displayed similar levels of replicative stress biomarkers, suggesting other contributing factors. We speculated that these subtypes might diverge in their growth profiles. To investigate this further, we inferred the duration and respective phases of their cell cycles from the normalised expression of the four classes of cyclins (details in **Online Methods**). Our analysis revealed that PACA 1 and 2 tumours exhibit comparable levels of cyclin D1 expression, which is higher than that of the PACA 3 subtype (**Extended Data Fig. 5h**). Moreover, cyclin D1 expression correlated with mutation burden at origins (*Rho* = 0.404, **Extended Data Fig. 5i**) indicating that cell proliferation is associated with mutational burden at origins. Additionally, we found that the PACA 1 subtype is characterised by a longer S-phase and shorter G1 and G2 phases compared to PACA 2 and 3 tumours (**Extended Data Fig. 5j**). Collectively, these observations demonstrate that increased cell proliferation and cell cycle dysregulation exacerbate the mutational burden at origins.

Our GSEA analysis also uncovered an association between the dysregulation of carbohydrate metabolism pathways and mutagenesis at origins (*P* = 9.84 × 10^-14^, permutation test, **Extended Data Fig. 5e**). To delve deeper into this observation, we examined the specific impact on glycolysis and the pentose phosphate pathway (PPP) in the PACA 1 tumour subtype (**Extended Data Fig. 5f**). Both pathways are intricately linked and exert significant influence on regulating DNA synthesis and repair in cancer. ^31^ Glucose-6-phosphate serves as a common substrate for both pathways, contributing to the production of pyruvate and ATP in glycolysis or ribonucleotides in the PPP. Inhibiting ribonucleotide synthesis is expected to affect DNA synthesis and repair. ^32^ Consequently, we hypothesised that the stimulation of glycolysis and inhibition of the PPP might elevate mutation rates at origins. Indeed, we observed a strong association (**Extended Data Fig. 5f**), with notable correlations between the expression levels of key enzymes and the ratio of mutation at origins versus flanks **Extended Data Fig. 5g**). For instance, both the expression of ATIC (a bifunctional enzyme involved in the final two steps of purine biosynthesis) and TALDO1 (an enzyme responsible for generating ribose-5-phosphate) exhibited a negative correlation with mutational burden at origins (*Rho* = −0.403 and −0.372 respectively). These findings suggest an unexpected interplay between metabolic pathways and mutagenesis at origins in cancers. Further investigations will be required to fully characterise this phenomenon.

### Constitutive origins are hotspots for genome rearrangements

Our previous GSEA findings (**Fig. 5d**) provide support for the downregulation of pathways associated with DNA replication and repair as a primary factor influencing the mutation rate at origins in pancreatic cancers (*P* = 3.73 × 10^-9^, permutation test, **Extended Data Fig. 5d**). To further explore this result, we dissected these pathways into smaller components and observed that the PACA 1 tumour subtype exhibits downregulation of cell cycle dependent DNA damage checkpoints and downregulation of factors involved in processing DNA single-strand breaks (**Fig. 6a**). Consequently, breaks originating at origins due to polymerase stalling at G4s are more likely to progress into S phase or mitosis without undergoing repair. This scenario is conducive to the emergence of genome rearrangements during cancer progression. ^33^ We then evaluated the prevalence of such events by focusing on structural and copy number variants, referred to as SVs and CNVs respectively. Our analysis revealed that PACA 1 and 2 subtypes are characterised by an increase in breakends at origins (**Fig. 6b**) and a higher number of copy number segments (**Fig. 6c**) compared to the PACA 3 subtype. These findings suggest that origins may serve as hotspots for genome rearrangement in pancreatic cancers.

**Figure 6.**
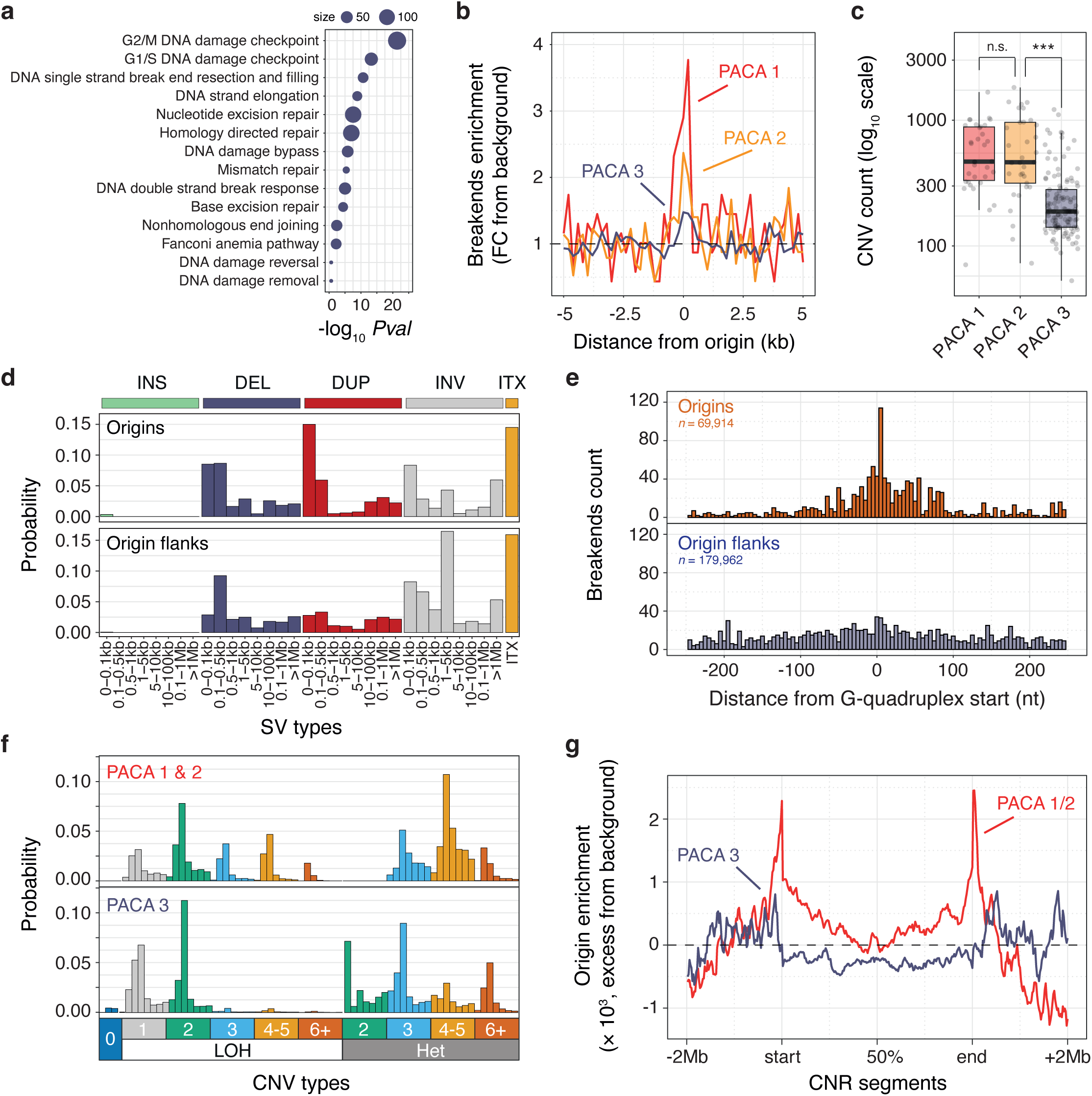
Constitutive origins serve as focal points for structural and copy number variants. **a**, Analysis of DNA repair and replication pathways. Primary KEGG pathways were dissected into smaller components to identify specific processes downregulated in PACA subtype 1 compared to subtypes 2 and 3. **b**, Distribution of structural variant (SV) breakends at origins across PACA tumour subtypes. Enrichment of breakends is presented as the fold change from background calculated from origin flank values. **c**, Distribution of the number of copy number variation (CNV) segments in individual PACA samples categorised by PACA tumour subtypes. Box plots illustrate medians and interquartile ranges, with grey dots indicating individual cancer samples. *n. s.* non-significant, \*\*\**P* < 0.001, Kolmogorov-Smirnov test. **d**, Pan-cancer signatures of structural variation based on breakends mapped at origins or origin flanks, categorised by SV types and lengths. SV types are: INS (insertion), DEL (deletion), DUP (duplication), INV (inversion) and ITX (translocation). **e**, Distribution of SV breakends at 5 nt resolution at G-quadruplex forming sequences within origin or origin flank domains. **f**, Signatures of copy number variation for PACA tumour subtypes, classified by CNV types, copy number, and evidence of loss of heterozygosity (LOH) or not (Het). Each CNV type is further classified by increasing segment sizes (from left to right). Segment sizes were excluded from this plot to enhance clarity. Detailed information on segment sizes can be found in the **Online Methods** section. **g**, Enrichment of origins within amplified CNV segments in PACA subtypes. Enrichment values were evaluated by partitioning CNV segments and their flanking domains (± 2 Mb) into an equal number of windows, calculating the number of origins per segment and per window, and then determining the mean number of origins per window. This value was then adjusted by subtracting the mean values observed in segment flanking domains. The resulting origin enrichment value indicates the excess of origins within a given window.

Due to the limited number of SVs observed at origins in PACA cancers, we expanded our investigation to characterise structural variation at origins across various cancer types. Our analysis confirmed an increase in breakends at origins pan-cancer (**Extended Data Fig. 6a**) and we proceeded with establishing the signatures of structural variation at origins and origin flanks using a standardised classification procedure. ^34^ This approach revealed an enrichment of short tandem duplications at origins compared to their flanks (**Fig. 6d**). These enriched duplication events occur to the detriment of other SVs (**Extended Data Fig. 6b**), exhibit a median length of approximately 100 base pairs (**Extended Data Fig. 6c**) and stem from breakends occurring predominantly at Gs (**Extended Data Fig. 6d**). This latter observation underscores the potential role of breaks originating from polymerase stalling at G4s as the primary driver of these events. To validate this hypothesis, we examined the distribution of SV breakends in the vicinity of G4s (**Fig. 6e**) and found an enrichment of breakends within 5bp of the start of G4 motifs at origins but not at G4s within origin flanks. Taken together, these findings indicate that breaks arising from replication initiation are transformed into a diverse array of short tandem duplications, posing the potential to alter genome structure and function.

Finally, we characterised the copy number segments mapped within PACA subtype genomes. Notably, we observed that the increase in segment numbers in PACA 1 and 2 (**Fig. 6c**) coincides with the presence of shorter segments with median lengths of ∼ 2 Mb (**Extended Data Fig. 6e**), suggesting a potential association between origins and focal copy number alterations. Analysing the pattern of copy number alterations in PACA subtypes, focusing on length, absolute copy number, and potential loss of heterozygosity (LOH) following an established procedure, ^35^ yielded distinct signatures for PACA 1 and 2 when compared to PACA 3 (**Fig. 6f**). Firstly, PACA 1 and 2 subtypes are characterised by the presence of high copy number segments (3 and above) that underwent LOH, indicative of chromosomal duplication and segmental aneuploidy, ^35^ possibly resulting directly from the replicative stress experienced by these subtypes. More interestingly, these subtypes also exhibited an increase in short segments (1 to 3 Mb) amplified to 3 to 5 copy numbers without signs of LOH. To explore a potential role for constitutive origins in generating these segments, we mapped the distribution of origins within PACA 1 and 2 amplified segments with copy numbers higher or equal to 3 and found that origins marked their boundaries (**Fig. 6g**). A permutation test confirmed the statistically significant enrichment of origins at the boundaries of these segments (P ≤ 0.001 over 1,000 permutations, **Extended Data Fig. 6f**). Importantly, such enrichment was not observed for amplified segments from the PACA 3 subtypes (**Fig. 6g**) or for loss segments across any PACA subtypes (**Extended Data Fig. 6g**). To better understand the mechanism by which breaks induced at origins drive copy number alteration, we noted that origin enrichment at segment boundaries was confined to amplified segments of 3 or 4 copy numbers (**Extended Data Fig. 6h**) showing no signs of LOH (**Extended Data Fig. 6i**). Collectively, these observations support the notion of constitutive origins as hotspots for copy number alteration and suggest that breaks originating from replication initiation drive focal rearrangements through a specific mechanism discussed below.

## Discussion

Our analysis identifies and characterises biological processes associated with replication initiation that focus mutagenesis and genome rearrangements at thousands of constitutive origins. Notably, while the origin domains we examined cover less than 0.3% of the human genome, they can contribute up to 2.7% of the mutations in some cluster 2 cancer samples. Considering that constitutive origins are enriched within functional regions of the human genome, ^6^ such as promoters and exon/intron junctions, we anticipate that this focused mutagenesis contributes to the evolution and expression of cancer genomes. Furthermore, we found that origin activity leads to the emergence of numerous tandem duplication events and copy number alterations with potential functional consequences. For instance, we identified the duplication of short origin sequences within the first introns of the MYC and TYK2 proto-oncogenes. These events have the potential to alter the expression of these genes, as origin sequences are enriched for specific transcription factor binding sites and *cis*-regulatory elements. ^6,36^ We also mapped the amplification of megabase segments, marked with constitutive origins at their boundaries, containing pancreatic cancer associated oncogenes such as EIF3B or MTA1 that may drive specific cancer phenotypes. ^37,38^

While the mechanisms underlying some of the mutational processes we identified are clear, further investigations are necessary to understand the molecular basis for origin-associated genome rearrangements. Although we observed distinctive signatures in SVs and CNVs at origins, the origins of these events remain speculative. It is noteworthy that double-stranded DNA breaks with protruding ends, such as those that may arise following polymerase stalling at G4s near origins, are known to trigger small tandem duplications following overhang gap filling and end-joining (reviewed in ^39^). In this context, the size of tandem duplication events would be controlled by the location of the breaks and the site where DNA synthesis is initiated. Intriguingly, tandem duplications are confined within the 1-kilobase domains defined by replication origins, leading to duplication of origin sequences. The CNV events we identified are also associated with constitutive origins, suggesting a role for breaks induced by replication initiation in their formation. Due to the absence of LOH in the resulting amplified segments, we can only speculate that they may result from a mechanism such as fork stalling and template switching (FoSTeS). ^40^ In this mechanism, the decoupling between leading and lagging strand synthesis, caused by polymerase stalling at G4s, would trigger a switch to a region with microhomology, such as another replication origin or G4-rich domains, to restart DNA synthesis. Such events could occur multiple times before the replication fork returns to its original position to continue replication to the chromosome end, resulting in the amplification without LOH observed in our analysis.

Our findings are based on the comprehensive analysis of mutagenesis occurring at ‘constitutive’ origins that are conserved across several human cell types. ^41^ These origins are recurrent sites for replication initiation and accumulate mutations at each cell division. However, it is important to acknowledge the potential variability in the activity and efficiency of these origins across different cancer types. Indeed, certain cancers may inactivate some origins or exhibit variations in their efficiency compared to cultured cells. Additionally, the induction of oncogenes can activate dormant origins, further complicating the estimation of global mutation rates at these sites. ^4^ Consequently, our estimations may be subject to some degree to both underestimation and overestimation. Nevertheless, our study reveals compelling associations across pan-cancer and tissue-specific analyses, strongly advocating for replication initiation playing a critical role in shaping and focussing specific mutational processes at origin sites. Notably, we demonstrate that the mutagenic potential of replication origins is exacerbated by replicative stress associated with tumour progression. Furthermore, we show that the intrinsic biology of cancer subtypes significantly influences the intensity of mutagenesis observed at these sites. These collective observations underscore the integral role of replication origins as fundamental genetic elements regulating the mutagenesis that drives the development and progression of human cancers.

## Online Methods

### Cancer mutation and associated datasets

Somatic mutation data for cancer, including single nucleotide variants (SNVs, generally referred to as mutations in the manuscript), structural variants (SVs), and copy number variants (CNVs), were sourced from the International Cancer Genome Consortium (ICGC, release 28). SNVs and SVs for individual cancer samples were derived from the Broad variant calling pipeline, while CNVs for pancreatic ductal adenocarcinoma samples were obtained from the DKFZ/EMBL variant calling pipeline. These mutations were directly used for analysis. Only cancer samples that were whole-genome sequenced and had more than 5,000 mutations were considered for computing SNV mutation rates, or 50 mutations at origin domains for single base substitution mutation signature analysis. Transcriptomic profiles were generated from sequencing-based gene expression data of ICGC tumours, focusing only on protein-coding genes. Gene expression levels were quantified as transcripts per million (TPM) using raw read counts and exon lengths associated with all transcripts mapped to a given gene. Exon lengths were retrieved from the TxDb.Hsapiens.UCSC.hg19.knownGene annotation R/Bioconductor package. Throughout the manuscript we refer to cancer types using the ICGC cancer project nomenclature and their associated acronyms; BOCA: Bone cancer, BRCA: Breast cancer, BTCA: Biliary tract cancer, CLLE: Chronic lymphocytic leukemia, CMDI: Chronic myeloid disorders, EOPC: Early onset prostate cancer, ESAD: Esophageal adenocarcinoma, GACA: Gastric cancer, LAML: Leukemia, LICA: Liver cancer, LINC: Liver cancer, LIRI: Liver cancer, MALY: Malignant lymphoma, MELA: Skin cancer, ORCA: Oral cancer, OV: Ovarian cancer, PACA: Pancreatic cancer, PAEN: Pancreatic endocrine neoplasms, PBCA: Pediatric brain cancer, PRAD: Prostate Adenocarcinoma and RECA: Renal cell cancer.

### Constitutive origin dataset

The constitutive origins analysed in our study were identified using our high-resolution origin mapping technique, initiation site sequencing 2 (ini-seq 2), in the EJ30 bladder carcinoma cell line. ^41^ These origin coordinates, available from the Gene Expression Omnibus archive with the accession number GSE186675, initially mapped in the hg38 human genome assembly, were converted to the hg19 assembly using the *liftOver* tool from the rtracklayer R/Bioconductor package, ^42^ with chain files obtained from http://hgdownload.cse.ucsc.edu. Previous research has shown significant overlap between these origins and those identified by short nascent strand isolation coupled with next-generation sequencing (SNS-seq) in the same cell line, as well as with a database of ‘core’ origins identified by SNS-seq in 19 other human cell samples. ^43^ To ensure accurate comparison of mutation rates, we considered 10-kb flanking regions on either side of origin midpoints. Additionally, to avoid potential biases from replication origin clusters, we focused only on domains containing a single origin, resulting in analysis of 9,341 constitutive origins.

### Defining genomic regions and mutation densities

In our analysis, we contrast mutation rates and mutational mechanisms at origin domains with those at their flanking domains. We define origin domains as 1 kb regions centred on origin midpoints, and origin flanking domains as 20 kb regions also centred on origin midpoints but excluding the previously defined origin domains. We calculated mutation rates near origins using 100 bp windows, counting the number of relevant mutations and adjusting for the number of covered bases. To assess mutational burden at origins, we computed ratios of the length-normalised mutation counts at origin domains relative to those in flanking regions.

### Single base substitution mutational signature analysis

Evaluation and visualisation of substitution mutational patterns at constitutive origins were performed using the MutationalPatterns R/Bioconductor package. ^44^ We defined mutation count matrices by considering the frequencies of each of the 6 pyrimidine substitution, calculated in each of the possible 96 trinucleotide 5′ to 3′ context. To compute background-adjusted signatures at origins, we adjusted mutation count matrices obtained from origin domains based on their trinucleotide composition and to similar mutation count matrices obtained from origin flanking domains. Tumour sample clustering was performed using these adjusted signatures to differentiate samples according to mutation processes and cosine correlation similarities were used to quantify the closeness of origin-associated signatures. Genome-wide signatures were not corrected for local variation in trinucleotide composition. Origin-associated mutational signatures were compared to known signatures of somatic mutations in cancerous human tissues collected by the Catalogue of Somatic Mutations in Cancer (COSMIC, release v3.2) ^10^ and available from https://cancer.sanger.ac.uk/signatures/sbs. Exposures to both newly reported and COSMIC signatures were computed in 100 bp windows using the fit_to_signatures and fit_to_signatures_strict function of the MutationalPatterns package respectively. For COSMIC signature refitting experiments, only signatures contributing to at least 50 mutations within at least a window were considered.

### Nucleotide excision repair profiles at constitutive origins

Genome-wide NER sequencing (XR-seq) datasets of cyclobutane pyrimidine dimers (CPD) and pyrimidine-pyrimidone (6-4) photoproducts (6–4 PP) from ultraviolet-irradiated CSB/ERCC6 mutant NHF1 skin fibroblasts, ^13^ were retrieved from Gene Expression Omnibus (GSE67941). To quantify repair against DNase I hypersensitivity (DHS) at origins, ENCODE consortium data for the GM04504 skin fibroblast cell line were used. Sequencing data were formatted into bigwig files to report read counts at 50 nt resolution. These files were used to compute coverage matrices and generate heatmaps using the EnrichedHeatmap R/Bioconductor package. ^45^ Correlations between mutation counts for cluster 1 tumours, XR-seq, and DHS signals at origins were computed based on averaged signals over origin domains (origin midpoints ± 500 bp). NER strand bias around origins was evaluated by analysing strand-resolved XR-seq signals in 100 bp windows, corrected for dinucleotide base composition. As reported, the MC-062 and 64M-2 antibodies were used for CPD and 6–4 PP XR-seq data generation, respectively, with MC-062 targeting thymidine dimers and 64M-2 targeting UV-C irradiated DNA, in which 6–4 PP adducts are generated at CC, CT, and TC dinucleotides. ^46^ Thus CPD and 6-4 PP XR-seq data were adjusted for the frequencies of these dinucleotides.

### G-quadruplex forming propensity

To assess the likelihood of a sequence folding into a G-quadruplex (G4) structure and to quantify the predicted stability of the resulting motif, we employed the G4H score, which is derived from the *G4 Hunter* algorithm. ^20^ This algorithm takes into account the richness and skewness of guanine (G) content within a sequence to compute its G4H score. Each position within a sequence receives a score ranging from −4 to 4. G-skewness is considered by assigning a neutral score (0) to adenine (A) and thymine (T), a positive score to guanine (G) (as it promotes G4 formation), and a negative score to cytosine (C) (as it promotes hairpin and impedes G4 formation). G-richness is accounted for by assigning a score equivalent to the length of G tracts for each G, with a maximum score of 4. Similarly, cytosines in C tracts receive negative scores. The G4H score for a sequence is the average of these individual scores. Thus, the propensity of a sequence to fold into a G4 motif correlates with its G4H score, with higher scores indicating G4 formation on the plus strand and lower scores on the minus strand. To investigate G4 formation at specific positions, we extracted sequence contexts (± 25 bp) and computed associated G4H scores. To identify potential G4-forming sequences, we searched for sequences conforming to the regular expression pattern N_5_G_3+_N_1–12_G_3+_N_1–12_G_3+_N_1–12_G_3+_N_5_ (where N represents any base), which represents a slight modification from the known *quadparser* algorithm, ^21^ allowing for extended loop lengths. Additionally, we expanded the core motif for G4 formation by including five flanking bases to account for the G4 context when predicting stability using the G4 Hunter algorithm.

### Differential gene expression and gene set enrichment analysis

Differential gene expression analysis was performed using the DESeq2 R/Bioconductor package, ^47^ with pancreatic ductal adenocarcinoma (PACA) samples grouped based on cancer subtypes identified through UMAP dimensionality reduction of their transcriptomic profiles. UMAP analysis was performed using the umap R package ^48^ from TPM values of protein coding gene expression using two components and starting from 20 random states. Changes in gene expression were computed by contrasting PACA subtype 1 with PACA subtypes 2 and 3 independently. Differentially expressed genes were ranked using the DESeq Wald statistic and subjected to gene set enrichment analyses (GSEA) using the fgsea R/Bioconductor package ^49^ and curated KEGG pathways. Hallmark gene sets sourced from the Molecular Signatures Database (MSigDB, version 7.1) ^50^ were used, available as R lists in RDS format at https://bioinf.wehi.edu.au/MSigDB/. Pathway enrichments were determined by comparing PACA subtype 1 with PACA subtypes 2 and 3 individually, followed by averaging normalised enrichment scores and combining *P* values using Fisher’s method. Pathways of interest were further dissected into smaller components using information from the Reactome pathway database (https://reactome.org/). ^51^ Once more, normalised enrichment scores and associated *P* values were combined from independent analyses comparing PACA subtype 1 with PACA subtypes 2 and 3. Consequently, our analysis delineates pathways that are specifically dysregulated in PACA subtype 1, which exhibit a higher mutational burden at origins.

### Replicative stress and cell cycle evaluation in pancreatic ductal adenocarcinoma

Replicative stress within PACA samples was evaluated from the gene expression profiles of established biomarkers. ^30^ Expression levels of these biomarkers were computed as Z scores relative to the distribution of raw count values across all PACA samples. Analysis of cell proliferation and cell cycle dysregulation in PACA samples was inferred from the expression patterns of the four cyclin classes: cyclin A (CCNA1 and CCNA2), cyclin B (CCNB1 and CCNB2), cyclin D (CCND1, CCND2, and CCND3), and cyclin E (CCNE1 and CCNE2) in transcripts per million (TPM). These expression profiles were used to estimate the cell population proportions in G1/S phase (calculated as the ratio of cyclin E expression to the sum expression of all cyclins), S/G2 phase (calculated as the ratio of cyclin A expression to the sum expression of all cyclins), and G2/M phase (calculated as the ratio of cyclin B expression to the sum expression of all cyclins). ^52^

### Signature of structural variation

Pan-cancer structural variant (SV) breakpoints were mapped to either origin or origin-flanking domains, and SV signatures within these domains were extracted following a standardised protocol. ^34^ SVs were annotated as INS (insertion), DEL (deletion), DUP (duplication), INV (inversion), and ITX (translocation) based on the orientation and position of the genomic location covered by the supporting reads, using a custom R script. Additionally, SVs were further classified based on their length into eight predefined intervals: 0-0.1 kb, 0.1-0.5 kb, 0.5-1 kb, 1-5 kb, 5-10 kb, 10-100 kb, 0.1-1 Mb and greater than 1 Mb. Finally, the frequency of occurrence for each SV event was computed.

### Signature of copy number alteration in pancreatic ductal adenocarcinoma and relationship with constitutive origins

To identify signatures of copy number alteration in PACA subtypes, we adapted an established procedure ^34^ with slight modifications to emphasise focal amplification of small segments. We combined segments from PACA subtypes 1 and 2 as they did not show significant differences in a preliminary analysis. Annotations from the ICGC data describing mutation types were used to evaluate loss of heterozygosity (LOH) or heterozygosity (Het) within CNV segments. Subsequently, segments were categorised based on their absolute copy number and length into seven predefined intervals: 0-0.1 Mb, 0.1-1 Mb, 1-2 Mb, 2-3 Mb, 3-5 Mb, 5-10 Mb, and greater than 10 Mb. The frequency of occurrence for each copy number variation (CNV) event was determined. We investigated the association between constitutive origins and copy number alterations by examining the distributions of origins within copy number segments. We first computed the number of origins per 100 kb window across the human genome, generating a bigwig file for subsequent coverage calculation at CNV segments. We then evaluated origin distribution by dividing CNV segments and their adjacent domains (± 2 Mb) into 100 equally sized windows and determining origin coverage per segment and per window using the normalizeToMatrix function of the EnrichedHeatmap R/Bioconductor package. ^45^ Enrichment values were then computed by averaging origin counts across windows within segments of interest. These values were adjusted for background by subtracting the mean counts observed in flanking domain segments. Consequently, the reported enrichment values represent the excess of origins compared to background levels. CNV segments with length greater than 100 kb and smaller than 5 Mb were considered for these analyses. To assess origin enrichment at the boundaries of amplified segments in PACA 1 and 2 cancer subtypes, we identified segment breakpoints as the start or end of mapped amplified segments. We evaluated the significance of overlap with origin domains using the *overlapPermTest* function of the regioneR R/Bioconductor package, ^53^ conducting 1,000 permutations.

## Computational and statistical analyses

Analysis and all statistical calculations were performed in R (version 4.0.3).

## Acknowledgements

We thank I. Clayson, T. Darling and J. Grimmet in Scientific Computing at the LMB for support.

## Contribution

The project was conceived by P.M. and J.E.S. P.M. was responsible for conceptualisation, methodology and formal analysis. P.M., G.G. and J.E.S. contributed to data interpretation. P.M. was responsible for writing the original draft. P.M., G.G. and J.E.S. reviewed and edited the manuscript.

## Extended Data Figure legends

**Extended Data Figure 1.**
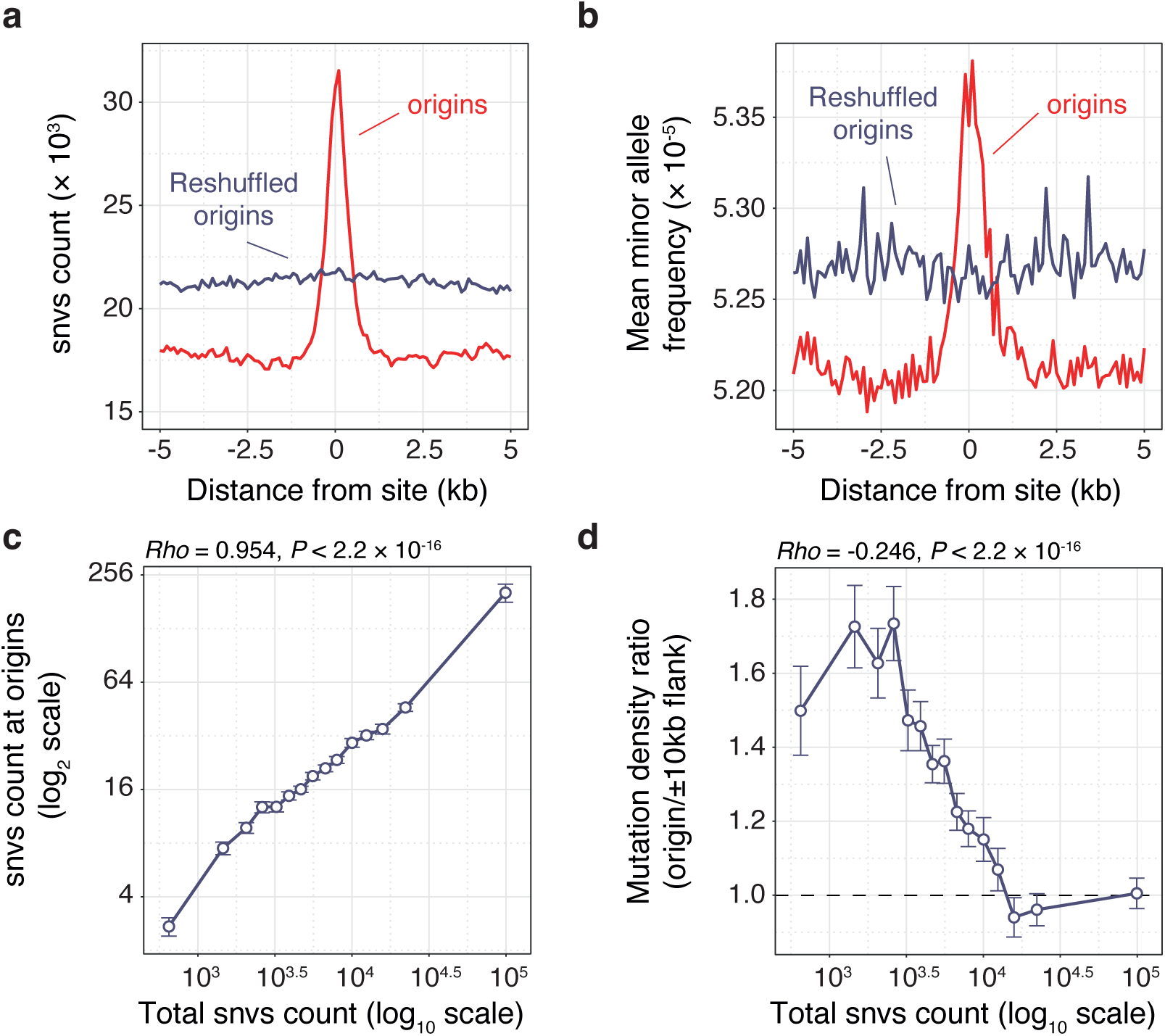
Mutational burden at constitutive origins. **a**, Count of single nucleotide variants (SNVs) and (**b**) mean allele frequency of alternative alleles, computed in 100 nt windows, at constitutive origins (red line) compared to control genomic locations obtained through the reshuffling of origin coordinates (blue line), from aggregated pan-cancer ICGC mutation data. **c**, SNV count at origins and (**d**) the ratio of mutation counts within origin domains over those in adjacent flanking regions relative to the total (genome-wide) SNV count for cancer samples with more than 5,000 called mutations.

**Extended Data Figure 2.**
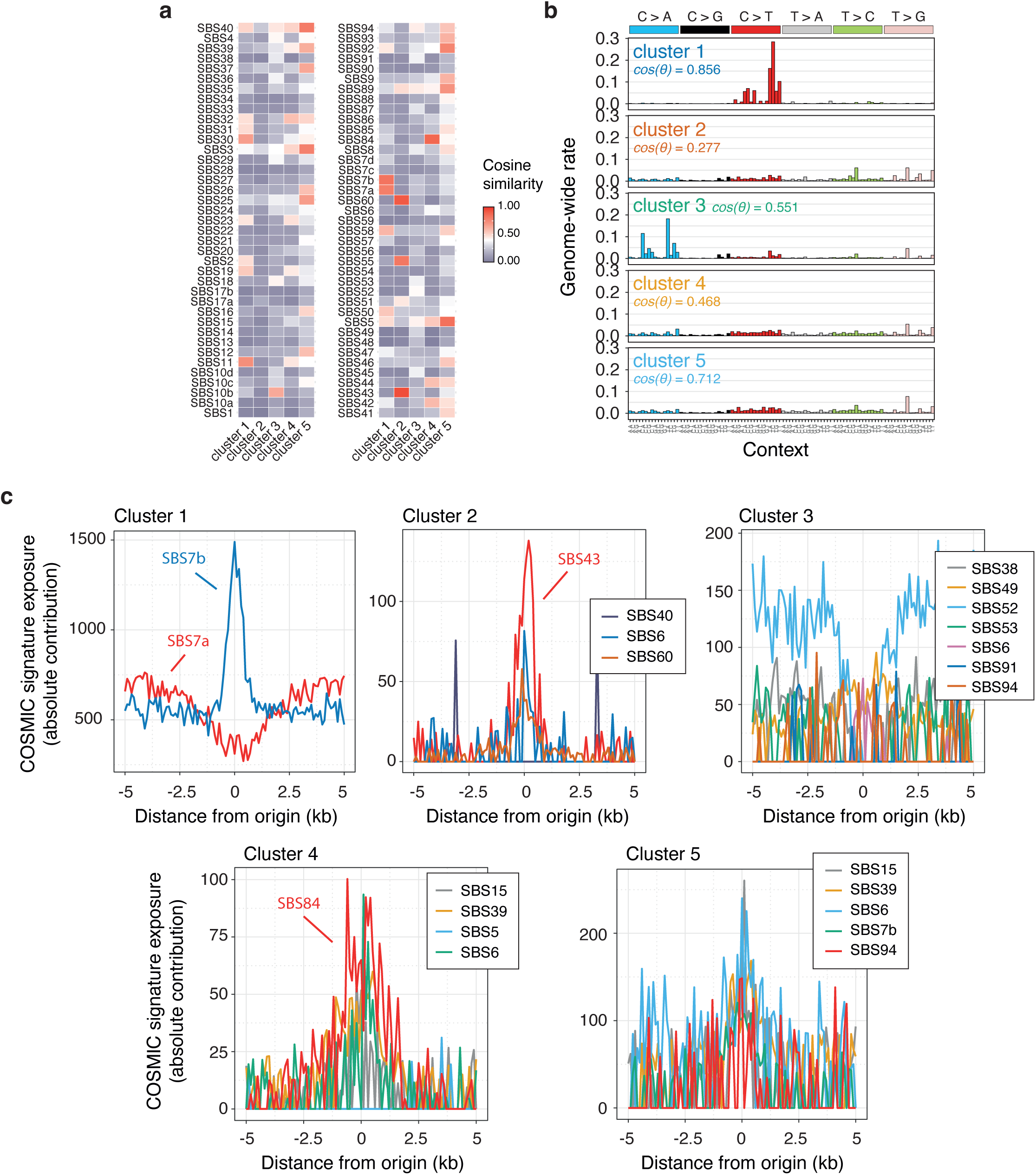
Characterisation of the mutational processes occurring at constitutive origins. **a**, Comparative analysis of origin-associated mutational signatures from tumours grouped into clusters 1 to 5, as defined in Fig. 2a, with established signatures of somatic mutations in human cancer tissues compiled by the Catalogue of Somatic Mutations in Cancer (COSMIC, release v3.2). The similarity between signatures is evaluated using cosine similarities. **b**, Genome-wide mutational signatures associated with tumours from clusters 1 to 5. Apart from cluster 1, origin-associated signatures display minimal resemblance to their respective genome-wide signatures, as indicated by the reported cosine similarities. **c**, Absolute contribution of COSMIC signatures to mutagenesis at origins, computed in 100 nt windows, for tumours in clusters 1 to 5. Signature contributions were computed using an iterative fitting procedure to exclude signatures with minimal contribution. Only signatures that contribute to more than 50 mutations within at least one 100 nt window are reported.

**Extended Data Figure 3.**
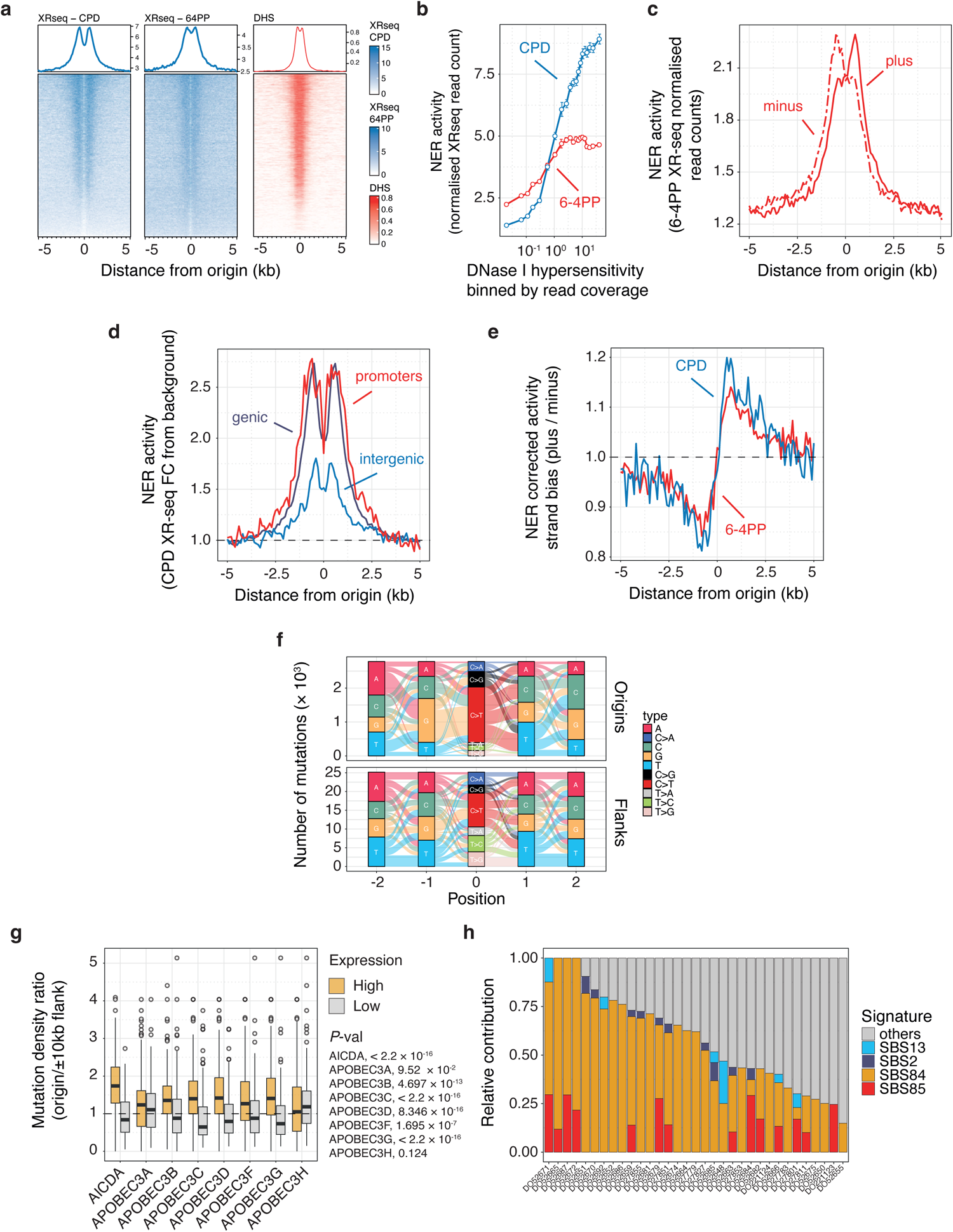
NER deficiency and deaminase activity at constitutive origins. **a**, Heatmaps displaying the coverage of XR-seq signals for CPD and 6-4 PP adducts (depicted in blues) and DNase I hypersensitivity (DHS, shown in reds) at constitutive origins. The heatmaps are arranged based on decreasing CPD XR-seq signal values. CPD and 6-4 PP XR-seq signals are derived from ultraviolet-irradiated CSB/ERCC6 mutant NHF1 skin fibroblasts, while the DHS signal is from the GM04504 skin fibroblast cell line. **b**, Evaluation of NER activity, quantified by the normalised read count of XR-seq signals for CPD and 6-4 PP adducts, in relation to origin accessibility, assessed by DHS read coverage, at origin domains (origin midpoints ± 1 kb). DHS signal is binned by read coverage, and NER activity represents the mean and standard errors to the mean of XR-seq read counts for each DHS bin. **c**, Strand-resolved XR-seq profiles for 6-4 PP at constitutive origins. **d**, XR-seq profiles for CPD at constitutive origins stratified by genomic location. **e**, NER-corrected activity strand bias computed from strand-resolved XR-seq signals for CPD and 6-4 PP in 100 bp windows. XR-seq signals were adjusted for dinucleotide base composition (refer to Online Methods). **f**, River plot illustrating the expanded contexts of mutations mapped within origin or origin-flanking domains for tumours of cluster 4. **g**, Mutation density ratios for individual malignant lymphoma (MALY) samples, organised by the level of AICDA (AID) or APOBEC3s activity. High (depicted by orange boxes) and low (depicted by grey boxes) expression levels correspond to the higher and lower tertiles of gene expression, respectively. Box plots depict medians and interquartile ranges, with individual cancer samples represented as circles. The reported *P* values compare values for high versus low expression levels using Kolmogorov-Smirnov tests. **h**, Relative contribution of COSMIC signatures on mutagenesis within origin domains of MALY samples with over 50 SNVs at origin domains. Specifically, SBS2 and SBS13 are associated to the activity of the APOBEC family of cytidine deaminases, while SBS84 and SBS85 are associated with the direct or indirect activity of AID. This analysis shows that AID is the main deaminase operating at constitutive origins.

**Extended Data Figure 4.**
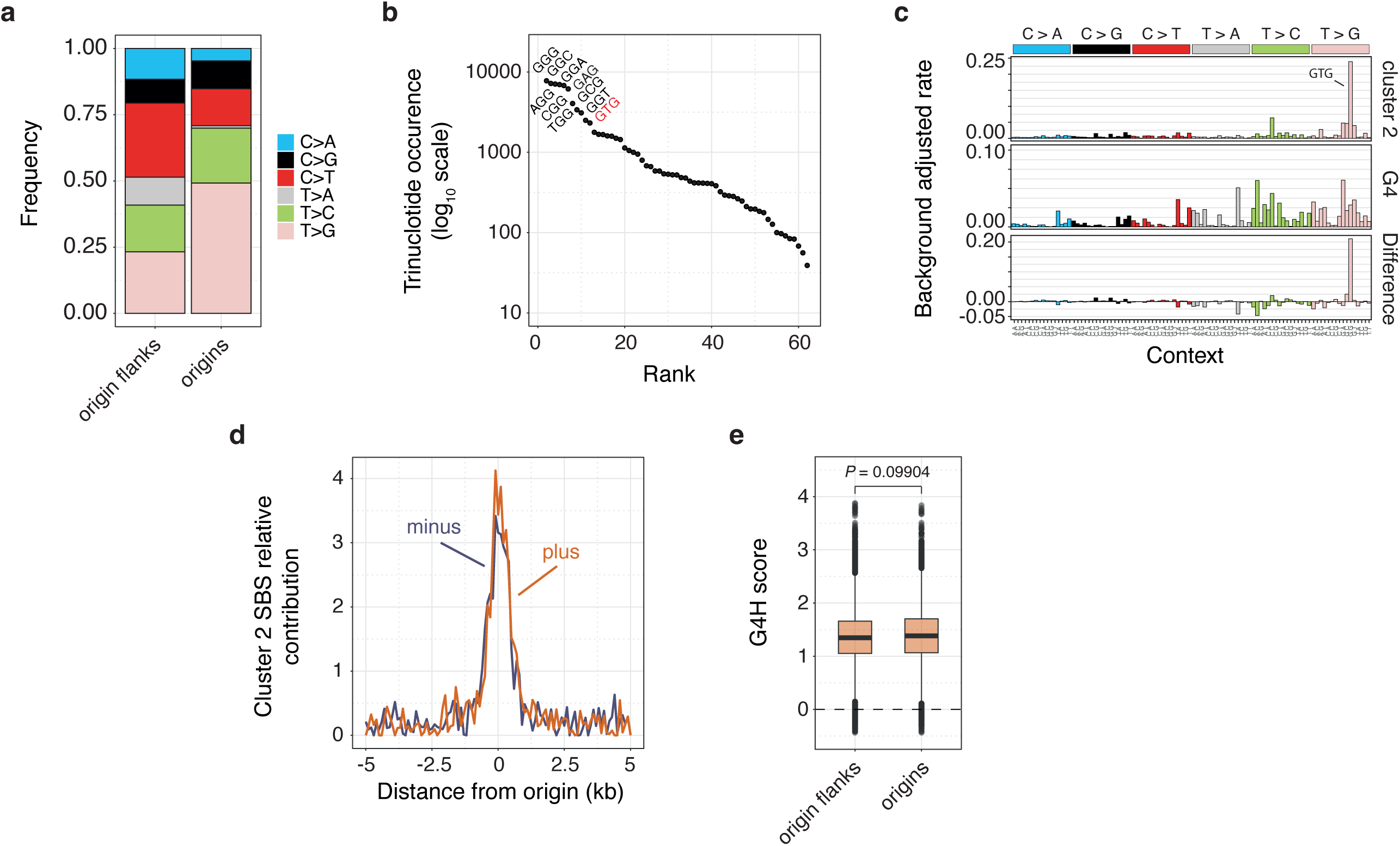
G-quadruplex mutational signature. **a**) Frequency distribution of each of the 6 pyrimidine substitutions within origin or origin-flanking domains of tumours from cluster 2, as defined in Fig. 2a. **b**) Trinucleotide occurrences within G-quadruplex (G4) forming sequences conforming to the pattern N_5_G_3+_N_1–12_G_3+_N_1–12_G_3+_N_1–12_G_3+_N_5_ (where N represents any base). The ten most recurrent trinucleotides are presented, with the characteristic GTG context highlighted in red. **c**, Background-adjusted mutational signatures extracted from origin mutations of cluster 2 or computed at G4 forming sequences corrected for G4 trinucleotide composition. The difference spectra underscore the overrepresentation of the GTG context. **d**, Strand-resolved contribution of cluster 2 mutational signatures to mutagenesis at the origins of cluster 2 tumours. Signature contribution was calculated by aggregating mutation calls from cluster 2 tumours. **e**, Distribution of G4 propensity scores, G4H scores, for G4 forming sequences conforming to the previous pattern found at origin or origin-flanking domains. Box plots depict medians and interquartile ranges, with outliers shown as grey dots. *P* value obtained from a Kolmogorov-Smirnov test.

**Extended Data Figure 5.**
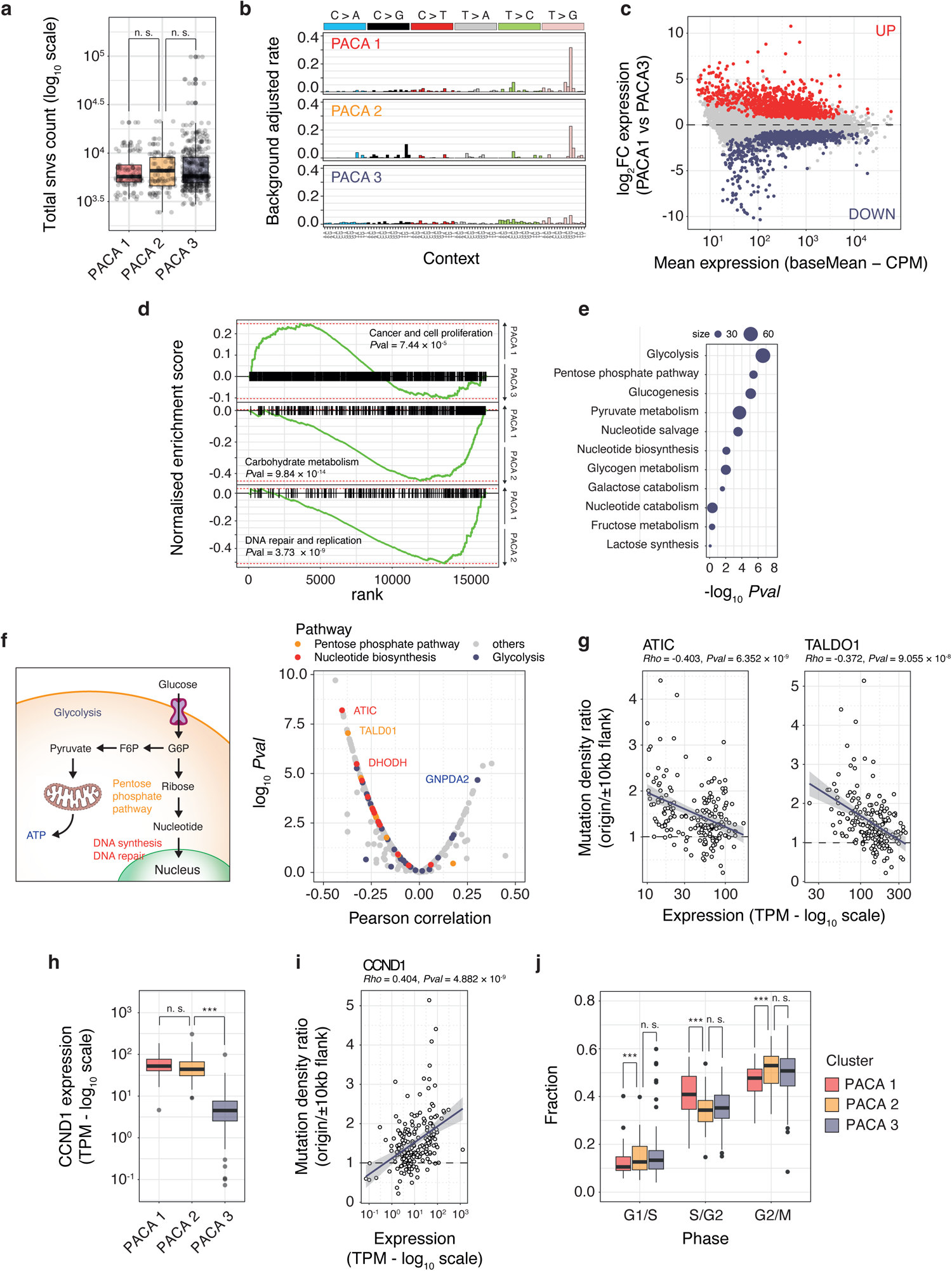
Determinants of origin mutagenesis. **a**, Distribution of genome-wide SNV counts for individual cancer samples categorised by PACA subtypes, as defined in Fig. 5a. **b**, Background adjusted mutational signatures at constitutive origin domains of PACA subtypes. **c**, DESeq2 differential gene expression analysis result contrasting gene expression in PACA subtype 1 versus 3. The plot reports differences in gene expression, expressed in fold change, in function of basal level of gene expression. The top 1,000 genes up- and down-regulated genes are reported in red and blue respectively. **d**, Gene set enrichment analysis associated with the primary pathways depicted in Fig. 5d. Enrichment scores were computed based on compiled gene lists related to Cancer and cell proliferation (top), Carbohydrate metabolism (middle), and DNA repair and replication (bottom). *P* values were derived by combining the *P* values associated with sub-pathways obtained from permutation tests and using Fisher’s method. **e**, Examination of Carbohydrate metabolism pathways. Primary KEGG pathways were dissected into smaller components to pinpoint specific processes exhibiting downregulation in PACA subtype 1 compared to subtypes 2 and 3. This analysis uncovered the dysregulation of Glycolysis and the Pentose Phosphate Pathway (PPP) in cluster 1 of PACA tumours. **f**. Schematic depiction (left panel) of the interconnection between the Glycolysis and PPP pathways. Glucose-6-phosphate acts as a common substrate for both pathways, contributing to pyruvate and ATP production in the Glycolysis pathway or ribonucleotide synthesis in the PPP. Thus, dysregulation of the PPP is anticipated to impact DNA synthesis and repair, resulting in increased mutagenesis at constitutive origins. To examine this hypothesis, we evaluated the Pearson correlation between the mutation ratio at origins versus origin flanks and the gene expression of enzymes involved in Carbohydrate metabolism (right panel). Enzymes associated with Glycolysis (blue), the PPP (orange), and Nucleotide biosynthesis (red) are highlighted. This analysis indicates that the expression of enzymes involved in the PPP and Nucleotide biosynthesis tends to negatively correlate with mutational burden at origins, underscoring a connection between glucose metabolism and mutagenesis at origins in pancreatic cancers. **g**, Negative correlations between the expression of key enzymes and mutational burden at origins is exemplified. ATIC is a bifunctional enzyme participating in the final two steps of purine biosynthesis, while TALDO1 is an enzyme responsible for generating ribose-5-phosphate. **h**, Expression levels of cyclin D1 in PACA tumours categorised in TPM. **i**, Cyclin D1 expression in TPM is function of the density ratio of mutations at origins to origin flanks for PACA samples. This analysis underscores the correlation between cell proliferation and mutational burden at origins. **j**, Fraction of cells in the G1/S, S/G2, and G2/M phases of the cell cycle for PACA tumours classified by subtypes. Cell cycle fractionation was inferred from the expression profiles of the four cyclin classes (refer to **Online Methods**): G1/S phase (computed as the ratio of cyclin E expression to the total cyclin expression), S/G2 phase (computed as the ratio of cyclin A expression to the total cyclin expression), and G2/M phase (computed as the ratio of cyclin B expression to the total cyclin expression). In panels **a**, **h**, and **j**, box plots illustrate medians and interquartile ranges, while individual cancer samples are depicted as grey dots. *n. s.* non-significant, \*\*\**P* < 0.001, Kolmogorov-Smirnov test.

**Extended Data Figure 6.**
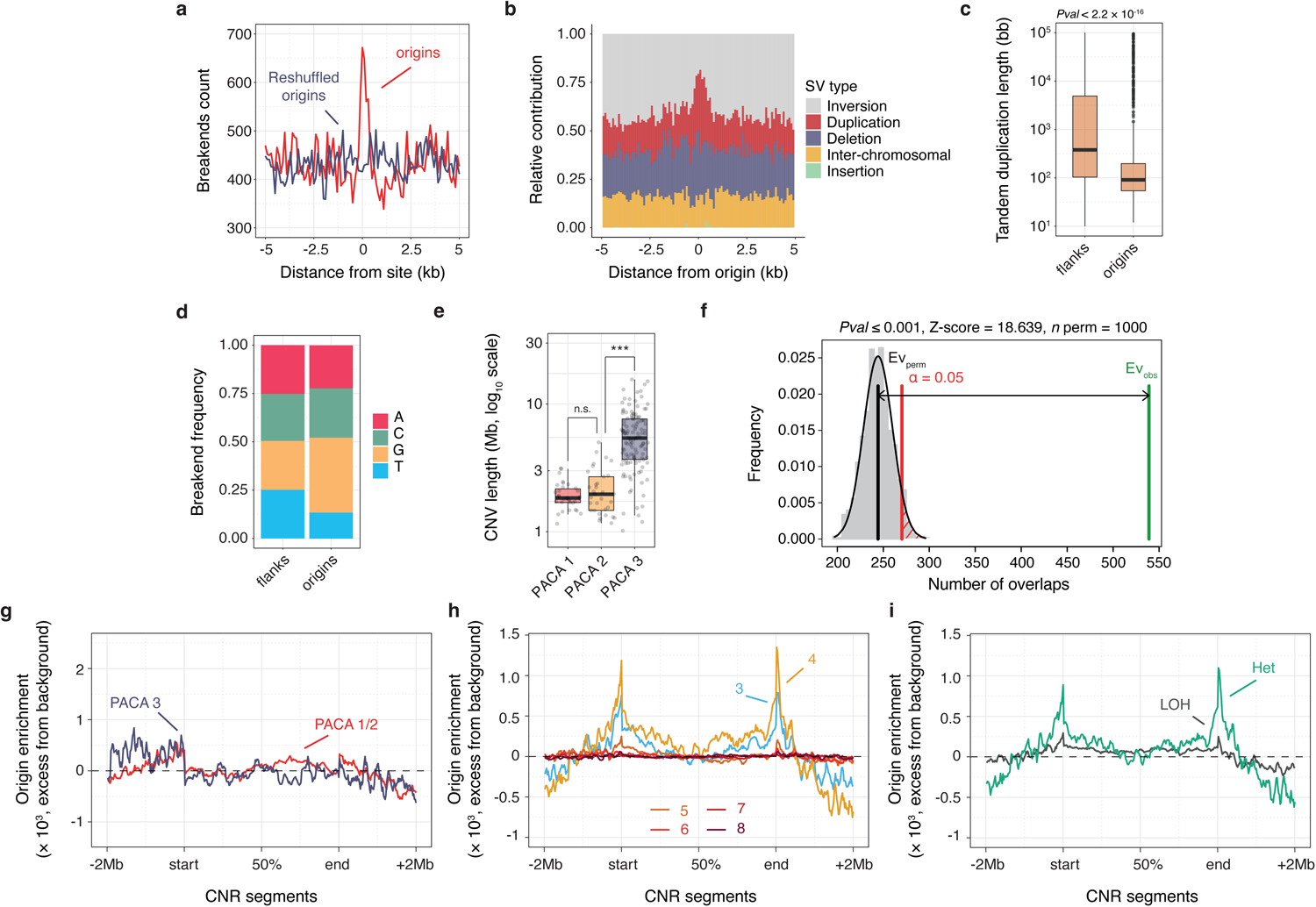
Constitutive origins are hotspots for genome rearrangements. **a**, Structural variants (SVs) breakends counts, computed in 100 nt windows, at constitutive origins (red line) compared to control genomic locations obtained through the reshuffling of origin coordinates (blue line), using aggregated pan-cancer ICGC mutation data. **b**, Relative contribution of SV type at constitutive origins. **c**, Distribution of the length of tandem duplication events mapped at origin or origin flanking domains. Box plots present medians and interquartile ranges. The *P* value is derived from a Kolmogorov-Smirnov test. **d**, Nucleotide frequency at SV breakends mapped at origin or origin flanking domains. **e**, Distribution of copy number variants (CNVs) length for individual cancer samples categorised by PACA subtypes, as defined in Fig. 5a. Box plots report medians and interquartile ranges, with individual cancer samples depicted as grey dots. *n. s.* non-significant, \*\*\**P* < 0.001, Kolmogorov-Smirnov test. **f**, Permutation test result assessing the enrichment of CNV breakpoints at constitutive origin domains. 1,000 permutations were performed, and the plot reports the distribution of randomised and observed numbers of overlaps. Enrichment of origins within CNV segments mapped in PACA subtypes considering (**g**) copy neutral and loss segments (copy number ≤ 2), (**h**) amplified CNV segments of different copy numbers and (**i**) amplified CNV segments of 4 copy numbers displaying either sign of loss of heterozygosity (LOH) or not (Het). Enrichment values were evaluated by partitioning CNV segments and their flanking domains (± 2 Mb) into an equal number of windows, calculating the number of origins per segment and per window, and then determining the mean number of origins per window. This value was then adjusted by subtracting the mean values observed in segment flanking domains. The resulting origin enrichment value indicates the excess of origins within a given window.

